# Viral adsorption to Moore swabs in passive wastewater sampling

**DOI:** 10.1101/2025.05.30.656954

**Authors:** Gouthami Rao, Timothy Purvis, Gyuhyon Cha, Jack Dalton, Michael Fisher, Katherine E. Graham, Konstantinos T. Konstantinidis, Yarrow Linden, Aaron Bivins, Margo A. Brinton, Christine Stauber, Joe Brown

## Abstract

Moore swabs have been used extensively for passive sampling in wastewater surveillance, typically yielding presence/absence information for targets of interest. Quantitative analysis of Moore swab data is only possible if target uptake is well characterized, specifically the relationship between quantity of the target in the liquid sample matrix and the quantity of target sorbing to the Moore swab as a function of time. The mechanism of Moore swab absorption remains unclear and is important to understand toward using them more quantitatively. We conducted viral adsorption and desorption experiments using nonpathogenic SARS-CoV-2 surrogates: Φ6, MHV, and BCoV as well as heat-inactivated Zika virus (ZIKV). We fit empirical adsorption data from batch experiments to Langmuir, Freundlich and Redlich-Peterson isotherm models. We observed the adsorption behavior of viral targets onto Moore swabs is best characterized by the Redlich-Peterson isotherm model. Moore swabs retained the highest viral RNA concentrations after exposure durations between 9-12 hours in the presence of target microbes during kinetic viral adsorption experiments. The results inform current and future use of Moore swabs to produce quantitative data during wastewater surveillance, especially in settings where composite sampling remains infeasible.

**Importance:** This paper describes the adsorption behavior of viruses and bacteriophages to Moore swabs. Passive sampling via Moore swabs is among the most scalable form of passive wastewater sampling, considered critical to advance wastewater surveillance globally. But key unknowns constrain the utility of Moore swabs and all passive sampling approaches, including the quantitative relationship between targets in wastewater and recovery via Moore swabs. Practical questions such as how long they should be deployed and whether they can be interpreted quantitatively really depend on a characterization of viral target loading behaviors on Moore swab material as a function of time and concentration in the wastewater. Here, we use an approach that is seldom applied to microbial targets to examine adsorption behavior of viruses to Moore swabs, deriving isotherms that describe the relationships between concentration of the viral targets in wastewater and time on attachment to swab material. This is a critical step in advancing the application of Moore swab passive sampling for wastewater surveillance, with potential relevance to other microbial targets of interest.

## Introduction

Wastewater monitoring includes both active and passive sampling methods. Active sampling refers to the collection of a sample through a grab or composite method by a person or autosampler at one or more specific time points. Passive sampling approaches involve deployment of sorption material in contact with the matrix to be left and sampled over a period of time (Vrana et al., 2005). Passive sampling methods have the potential advantages of being cost-efficient and easily deployable (Habtewold et al., 2022), but also come with limitations including poorly characterized sorption capacity and kinetics (Yang et al., 2022).

The application of passive sampling devices for wastewater surveillance has expanded beyond the original Moore swab method, first implemented to identify chronic carriers of *Salmonella enterica* serotype typhi and *Salmonella enterica* serotype paratyphi during a 1946-1949 typhoid outbreak in sewage drains of urban Sidmouth, England (Moore, 1951, 1948). Once microbial detection using Moore swabs was demonstrated to be useful, additional targets in wastewater such as poliovirus (Dániel and Dömök, 1962), *Vibrio cholerae* (Barrett et al., 1980), and *Aeromonas* spp. (Villarruel-López et al., 2005) were also investigated. The method also garnered attention in Santiago, Chile to identify *Salmonella enterica* serotype typhi in crop irrigation waters (Sears et al., 1984). In addition to irrigation water and wastewater, Moore swabs have also been used to detect pathogens in surface waters (Sikorski and Levine, 2020) and most recently for building-level wastewater assessment on university campuses for the detection of SARS-CoV-2, the etiological agent for COVID-19 (Gibas et al., 2021; Liu et al., 2022; Scott et al., 2021).

Because of the potential advantages of using Moore swabs during the COVID-19 pandemic (Liu et al., 2022; Schang et al., 2021), passive sampling methods have recieved renewed attention and innovation. Novel passive sampling methods included 3D-printed “torpedoes” housing cotton gauze, an electronegative membrane, and cotton bud all within a plastic housing shaped like a torpedo to avoid ragging (i.e., agglomeration of solids or materials that catch and stick to the sampler) (Schang et al., 2021). Other methods included a porous, ball-like structure from Canada that also housed electronegative membranes (Hayes et al., 2021), commercially-available tampons (Bivins et al., 2022b), and various kinds of Moore swabs for surveillance via suspension in wastewater flows (Cha et al., 2023; Liu et al., 2022).

Previous work using Moore swabs, or cotton gauze pads, often focused on the quantal (presence-absence) uptake of a specific pathogen target and performance often equaled or exceeded other sampling methods, such as grab or manually retrieved composite samples, especially in upstream settings with intermittent flow (Habtewold et al., 2022; Hayes et al., 2021; Rafiee et al., 2021; Schang et al., 2021). For example, Rafiee et al. determined that Moore swabs and composite samples were more sensitive to detect SARS-CoV-2 than grab samples from sewage manholes, and that Moore swabs even performed comparably to a 16-hour composite sample in terms of qualitative presence/absence and mean Cq values.

Despite the potential advantages, cost-effectiveness, and demonstrated utility of deploying Moore swabs as a passive sampling method, gaps remain about how quantitative data from them should be interpreted (Bivins et al., 2022a). Viral loading on Moore swabs has not been adequately characterized, nor has the quantitative relationship between viral titer in wastewater matrices and viral quantity recovered on the Moore swab recovery rates been determined.

Describing viral loading over time as a function of viral titer in the wastewater to be sampled – and, relatedly, desorption once the target virus is no longer present – is required to understand how to usefully interpret quantitative molecular data from this widely used and re-emerging passive sampling approach (Sikorski and Levine, 2020). To this end, we parameterized viral adsorption isotherms for viral loading onto Moore swabs under controlled experimental conditions.

Isotherms are generalizable quantitative models that can be used to estimate target loading for conditions reasonably, but not exactly, similar to the experimental conditions used to characterize the model. Three commonly used isotherm models are Langmuir, Freundlich, and Redlich-Peterson. Langmuir models assume monolayer adsorption and a relatively homogenous surface. Freundlich models assume non-uniform adsorption sites on heterogeneous surfaces, and Redlich-Peterson assumes both monolayer and multi-layer adsorption mechanisms (Allen et al., 2004). We determine how these isotherms perform based on model fitting, but also consider biological plausibility based on the specific assumptions associated with each isotherm type and hypothesize that complex adsorption mechanisms are at play with Moore swabs viral uptake and desorption. We further examine the relationship between viral loading over time as viral titer is held experimentally constant and describe desorption behavior once the target is no longer present.

## Materials and Methods

### Reagents and Materials

We used premium cotton gauze rolls purchased from the Mighty-X store (Clackamas, Oregon, USA) to construct the Moore swabs. Gauze was cut into 48-inch length x 4-inch width and folded in a Z-fold fashion six times. Each swab weighed an average of 5.0 grams. We secured folded swabs to a 50 lb. fishing line before placing them in a biological safety cabinet flow hood for UV treatment for 15 minutes on each side. We stored treated Moore swabs in sterile sample bags (VWR, 22 oz).

We conducted batch equilibrium and kinetic experiments to determine when forward and reverse reactions stabilize and characterize the sorption of viral targets over time. Batch and kinetic experimental set-ups were initially designed using previous work (Brown et al., 2001), with modifications for the continuously-stirred tank reactors (CSTR) in the kinetic and desorption experiments detailed further in Supplementary Text C. We selected pseudomonas bacteriophage (Φ6), murine hepatitis virus (MHV), and bovine coronavirus (BCoV) as nonpathogenic surrogates for SARS-CoV-2 and heat-inactivated Zika virus (ZIKV) as potential viral targets for wastewater surveillance efforts (Ahmed et al., 2020). Φ6 stock titers were pre-determined by double agar layer plaque assay, ZIKV stock titers were validated by immunofocus assay, and BCoV and MHV stock concentrations were determined using digital PCR (dPCR). Details on stock preparations are included in Supplemental Text A. Stocks were originally donated by Yarrow Linden, Dr. Michael Fisher, and Dr. Margo Brinton.

For equilibrium experiments, we collected 10L of final effluent and 5L of primary sludge from a local wastewater treatment facility in Chapel Hill, North Carolina on two separate occasions between July 2022 and February 2023. We transported all samples on ice and stored them at 4°C until experiments were conducted within one month. For kinetic experiments, we collected 5L of final effluent and 1L of primary sludge from the same wastewater treatment facility in Chapel-Hill, North Carolina on July 22, 2022. Final effluent was combined with primary sludge and autoclaved at 121°C for 15 minutes.

### Moore Swab Processing – skim milk flocculation

For each equilibrium and kinetic experiment, the processing of Moore swabs remained the same. We placed each Moore swab into a sterile potato ricer and pressed the sorbate into a sterile polyethylene beaker. Contents were poured into 50 mL centrifuge tubes with the total swab eluate recorded. We then used a skimmed milk flocculation method outlined elsewhere (Philo et al., 2021). We added 1 mL of a 5% skim milk solution per 100 mL of wastewater eluate and adjusted the skimmed-milk-eluate solution pH to 3.0—4.0 using 1M hydrochloric acid. We placed sample eluates on a platform shaker plate (Innova 2100, Marshall Scientific, Hampton, NH) at room temperature (20-25°C) at 200 revolutions per minute (rpm) for two hours. After shaking, we centrifuged samples at 3500 x g at 4°C for 30 minutes and the supernatant was discarded. The resulting pellet was archived at –80°C until batch extractions were complete.

### Pretreatment and RNA Extraction

We extracted each pellet using the QIAamp 96 Viral extraction kit on the QIAcube HT instrument (Qiagen, Hilden, Germany). We followed pretreatment steps outlined elsewhere that were previously optimized for fecal sludge (Capone et al., 2020). Briefly, we used 1 mL of ASL stool lysis buffer (Qiagen, Hilden, Germany) per 200 μL of pellet. We used the TissueLyzer II for 5 minutes at 25,000 Hertz and then rotated samples to face the opposite direction and repeated the step. We incubated samples for 10-15 minutes at room temperature before we centrifuged for 2 minutes at 14,000 rpm. We transferred 200 μL of the resulting supernatant into the S-block and used the automated QIAcube HT protocol. We included a negative extraction control for each batch. We stored the extracts at –80°C until samples were processed using a digital PCR platform.

### Digital PCR Processing

We processed all samples and stock concentrations using digital PCR (dPCR) on the QIAcuity Four (Qiagen, Hilden, Germany). Primer and probe sequences and references are presented in Supplementary Table A. Each reaction contained 10 μl of 4X QIAcuity One-Step Viral RT-PCR Master Mix, 0.4 μL multiplex 1X RT mix, 2 μL of template RNA, and molecular grade water for a total reaction volume of 42 μL. The final assay was performed as a four-plex after single-plex validation. All primer and probes were procured from Integrated DNA Technologies (IDT, Coralville, IA). Final forward and reverse primer concentrations for BCoV were 3200 nM and probe concentration of 400 nM TAMRA probe. Final forward and reverse primer concentrations for ZIKV were 800 nM and probe concentration of 200 nM SUN probe. Final forward and reverse primer concentrations for MHV were 800 nM and probe concentration of 400 nM ROX probe. Final forward and reverse primer concentrations for Φ6 were 3200 nM and probe concentration of 400 nM FAM probe. The reaction mixture was loaded onto QIAcuity Nanoplate 26k 24-well plates and run on a QIAGEN QIAcuity Four machine operated with the QIAcuity Software Suite 2.1.7.182 (Qiagen, Hilden, Germany). All final cycling conditions include: 50°C at 40 minutes for reverse-transcriptase step, one initial heat activation cycle at 95°C for 2 minutes, and 40 denaturation cycles at 95°C for 5 seconds and 50°C for 30 seconds for annealing/extension. Imaging steps were reduced to the following per channel: red at 210 m/s, yellow at 350 m/s, green at 400 m/s. The orange channel was run and imaged separately at 400 m/s. Thresholds (i.e., partition classification) were determined as 130 RFU (fluorescence intensity) for BCoV, 80 RFU ZIKV, 50 RFU Φ6, and 75 RFU MHV. We processed all samples in triplicate and included one no-template control and one positive control for each plate.

### Equilibrium Experiments

We assessed equilibrium saturation behavior of Moore Swabs by spiking various concentrations of Φ6 (10^8^ gene copies [gc]/mL, 10^7^ gc/mL, 10^6^ gc/mL), MHV (10^5^ gc/mL, 10^4^ gc/mL, 10^3^ gc/mL), BCoV (10^8^ gc/mL, 10^7^ gc/mL, 10^6^ gc/mL), and heat-inactivated ZIKV concentrations (10^5^ gc/mL, 10^4^ gc/mL, 10^3^ gc/mL) into each polyethylene, sealed container seeded with varying concentrations of wastewater effluent/solids and mixed at 200 RPM for 15 minutes. We placed individual Moore swab of three different masses (1g, 2g, and 5g) into each container. Each combination of surrogate concentration and swab mass was replicated in triplicate. The containers were continuously mixed via a shaker platform for 24 hours at a rate of 250 rpm. We processed all Moore swabs as outlined in section 3.2.2. immediately after collection. We tested wastewater concentrations at low, medium, and high TSS wastewater primary effluent + solids with a total of 100 mL per beaker. The TSS target ranges were 100-200 mg/L, 200-300 mg/L, and 400-500 mg/L, respectively (Hayes et al., 2022).

### CSTR set-up for kinetic experiments

We designed the continuously-stirred tank reactors (CSTRs) with the intention of creating a well-mixed matrix for Moore swab adsorption. Fundamentally, a CSTR assumes the targets are instantly and uniformly mixed throughout the reactor, meaning any sampling would be representative at any point in the reactor. The CSTRs had an optimal design volume for 10-12 L with a counterclockwise stirring direction that allowed for the recirculation to push to the bottom of the reactor, employ a homogenous mixing, and provide enough spacing so no two swabs touched. The wastewater matrix contained 20% primary effluent (v/v), 10-40 mL of primary sludge as our solids addition, and the remaining was dechlorinated tap water until a total volume of 10L was reached. We also used rechargeable batteries at 12V (Mighty-Max, Edison, NJ) to allow the CSTR motors to run for the duration of the experiment.

### Kinetic Experiments

Based on the equilibriums experiments, we determined the optimal viral concentration to test for kinetic experiments. Uptake behavior for viral targets was assessed by placing Moore swabs in a CSTR. Details on CSTR design parameters are included in Supplementary Text C. Each CSTR contained a 20% wastewater effluent at varying TSS conditions and was spiked with targets BCoV, Φ6, MHV, and ZIKV. Sampling of the swabs occurred over 5, 15, 30, 45, and 60 minutes and 3, 6, 9, 12, 24, and 48 hours. A process control of a CSTR containing all targets minus swabs in same wastewater effluent + TSS mixture was also tested at the same time points to account for degradation. The final bulk wastewater volume was 10L per reactor.

We conducted this experiment by measuring the Moore swab mass in a dry state with an aluminum weighing dish, then placing the swabs into the CSTRs at the indicated effluent concentrations and removed the swabs at the same time points for the kinetic experiments (5, 15, 30, 45, 60 minutes and 3, 6, 9, 12, 24, and 48 hours). Post-ricer-squeezed Moore swab samples were weighed on the weigh dish and total mass was measured. We placed samples in an oven at 105°C for 24 hours and total mass recorded after 24 hours. We ran samples in duplicate for each primary effluent + TSS concentration.

Each reactor contained six swabs and four reactors were used to complete one TSS category, including duplicate swabs. Once we finished sampling all relevant time points from one reactor then we disinfected the reactors with 10% sodium hypochlorite (v/v) for a contact time of at least 1 hour, disposed of down the drain, washed and autoclaved until next use.

### Desorption Experiments

Scour, or desorption, is defined as the physical process and release of molecules or particles from a surface. To assess desorption from Moore swabs, we first saturated Moore swabs with our targets for 24 hours on a shaker plate using the same spiked viral concentrations as the kinetic experiments. Scour behavior was assessed by submerging the saturated Moore swabs and placing in a CSTR of dechlorinated tap water. Moore swabs were pulled out of the system and sampled at 5, 15, 30, 45, and 60 minutes and 3, 6, 9, 12, 24, and 48 hours. Sampling was completed in triplicate for the first time points and in duplicate for the last time point. We also tested a process control of a CSTR containing Φ6, MHV, and ZIKV target in bulk water at the same time points to account for target degradation.

### TSS, pH, and temperature measurements

We conducted TSS measurements using ProWeigh 47 mm, borosilicate glass fiber filters (Environmental Express, Charleston, South Carolina) following the Standard Methods for the Examination of Water and Wastewater 2540D (“2540 SOLIDS,” 2017). We ran negative control blanks after every ten samples processed. We recorded temperature and pH at the beginning and end of each set of experiments using a low-range pH/electrical conductivity Hannah multi-meter (Smithfield, RI, USA). Kinetic experiments had temperature and pH checks at the 5 min, 12-hr, 24-hr, and 48-hr time points.

### Data Analysis

First, we took our measured variables such as the initial liquid phase viral concentration (*C*_0_ – gene copies/mL), our equilibrium liquid phase viral concentration (*C*_*e*_ – gene copies/mL), reactor volume (liters) and Moore swab masses (grams) to calculate our equilibrium viral concentration on the swab (*Q*_*e*_ – gene copies/gram) as seen in Equation 1.

Equation 1. Mass/Mass ratio of viral target to swab:

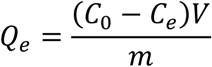

We considered the Langmuir, Freundlich, and Redlich-Peterson isotherm models (equations 2-4) to assess adsorption of viral RNA targets to the Moore swabs, which has been used in similar applications by others for metals or other adsorbent materials (Brown et al., 2001; Lasheen et al., 2012).

Equation 2. Langmuir isotherm

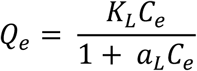

Equation 3. Freundlich isotherm

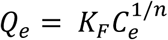

Equation 4. Redlich-Peterson

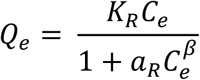

Then we generated linearized isotherm plots for each model type to determine isotherm constants (K_L_, K_F_, and K_R_) and correlation coefficients (a_L,_ *n*, and β) via model parameterization. Linearized expressions for each model are included as equations 5-8.

Equation 5. Langmuir (linearized)

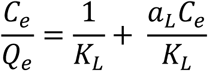

Equation 6. Freundlich (linearized)

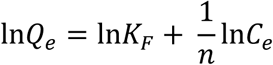

Equation 7. Redlich-Peterson (linearized)

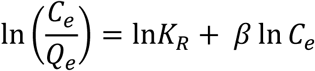

This information will help ascertain the rate of adsorption and desorption occurring on the Moore swabs and the models generated can be used to extrapolate a variety of concentrations. We used the data generated from these experiments to develop mass balance models to better understand target loading on the Moore swabs. Once the relationship between target concentration, time, and mass of Moore Swabs is identified any flow condition can be simulated.

R Studio version 4.2.1 ggplot2 package was used complete graphs and conduct linear regressions to obtain or calculate Langmuir (Langmuir, 1916), Freundlich (Freundlich, 1906), and Redlich-Peterson (Redlich and Peterson, 1959) constants and coefficients. The best fit estimates were based on R^2^ values > 0.8.

## Results

### TSS, pH and temperature summaries

The following results reflect the TSS, pH and temperature summary statistics for each experiment type. For batch equilibrium experiments, the average temperature was 68.5°F, with a pH of 7.25. Kinetic experiments had an average temperature of 65.7°F and pH of 6.2, and desorption experiments had an average temperature of 68.6°F and pH of 6.0. Table 1 includes a breakdown of average TSS and standard deviations, by the pre-determined categories.

**Table 1.**
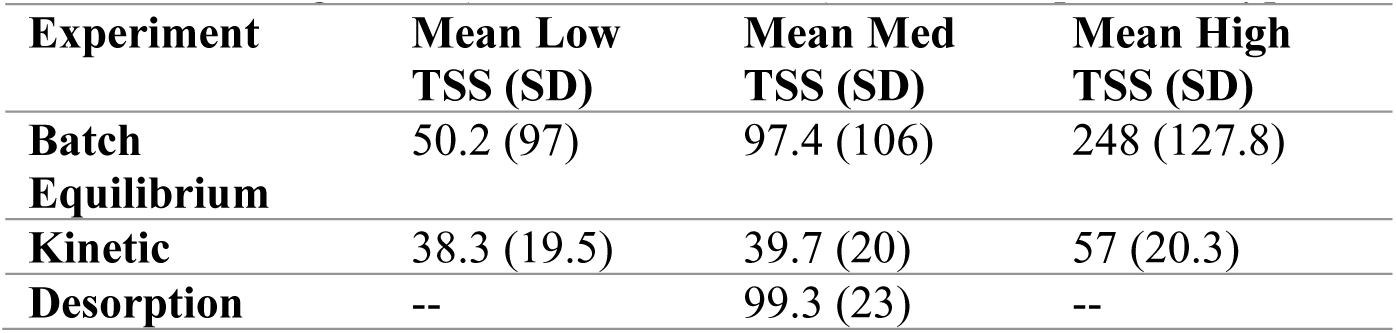
Average TSS (standard deviations) for each experiment type.

| Experiment | Mean Low<br>TSS (SD) | Mean Med<br>TSS (SD) | Mean High<br>TSS (SD) |
| --- | --- | --- | --- |
| Batch<br>Equilibrium | 50.2 (97) | 97.4 (106) | 248 (127.8) |
| Kinetic | 38.3 (19.5) | 39.7 (20) | 57 (20.3) |
| Desorption | -- | 99.3 (23) | -- |

### Equilibrium capacity by target and by TSS

Equilibrium experiments run for 24 hours indicated that maximum adsorption capacity was 1.1 x 10^13^, 4.5 x 10^11^, 3.5×10^12^, and 1 x 10^11^ gene copies per gram of swab for BCoV, MHV, Φ6, and ZIKV, respectively (Supplementary Figure D1). Regardless of TSS level stratification (Supplementary Figure D2), we observe an increase in equilibrium capacity as our equilibrium viral target concentration increases.

### Equilibrium Isotherm Models

We plotted the Langmuir and Freundlich isotherm models and due to poor fit (R^2^ value < 0.5) we included the data as Supplementary Figures E-H, which includes compiled data and data stratified by TSS categories to note linear directionality, if any. The Langmuir models showed poor fit to the data and resulted in a correspondingly low coefficient of determination (R^2^) values for BCoV (0.16), MHV (–0.01), Φ6 (–0.14), ZIKV (0.71), whereas the Freundlich models demonstrated similarly low R^2^ values with BCoV (0.08), MHV (0.0), Φ6 (0.78), and ZIKV (0.41). However, the Redlich-Peterson isotherm plots (R^2^ value = 0.7 – 0.88), overall and by TSS are presented in Figures 1 and 2, respectively. Table 2 depicts the Langmuir, Freundlich, and Redlich-Peterson constants and coefficients and further stratified by TSS.

**Fig. 1.**
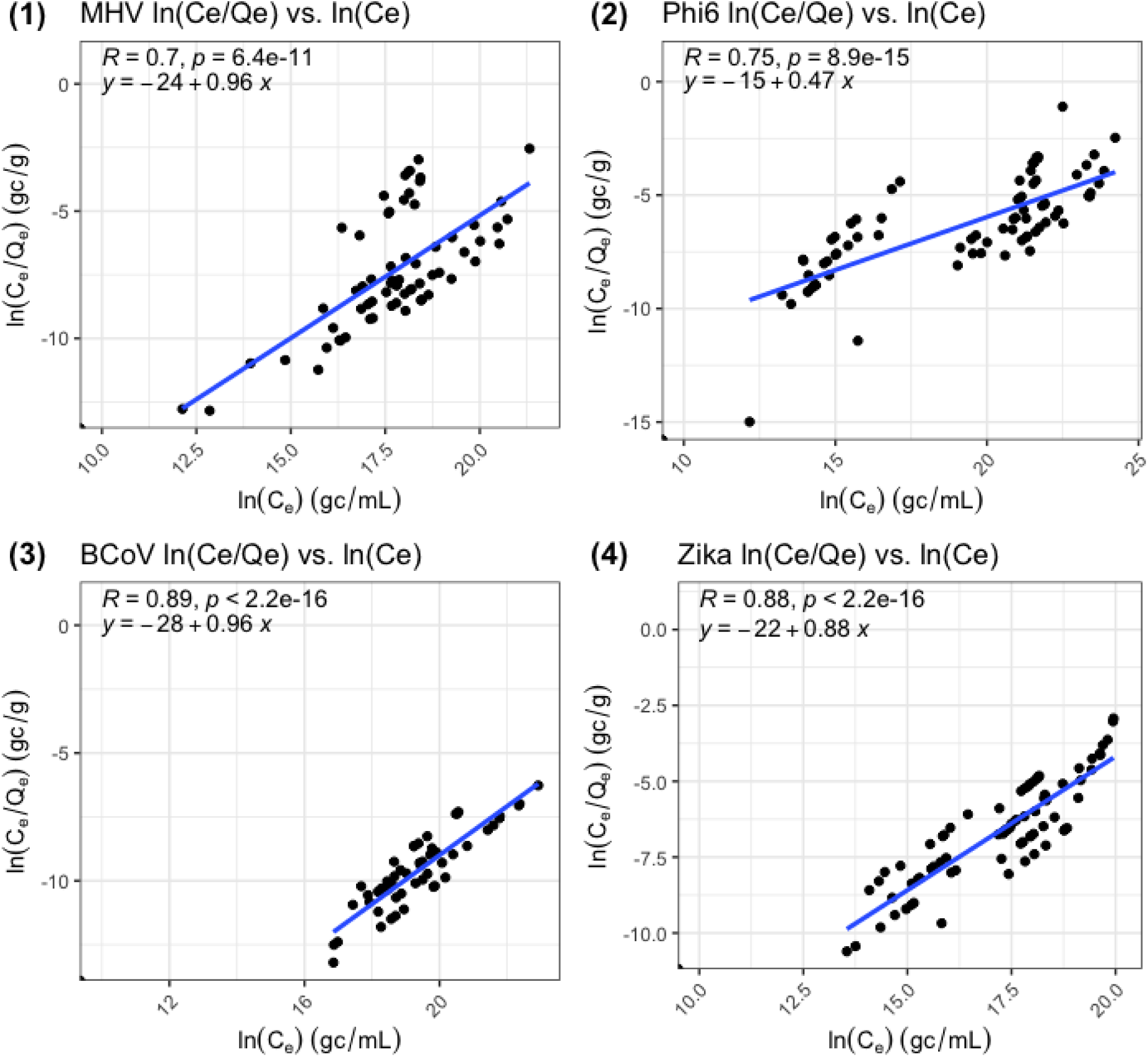
Final Redlich-Peterson isotherm plot for (1) MHV; (2) Φ6; (3) BCoV; (4) ZIKV. R = r^2^.

**Fig. 2.**
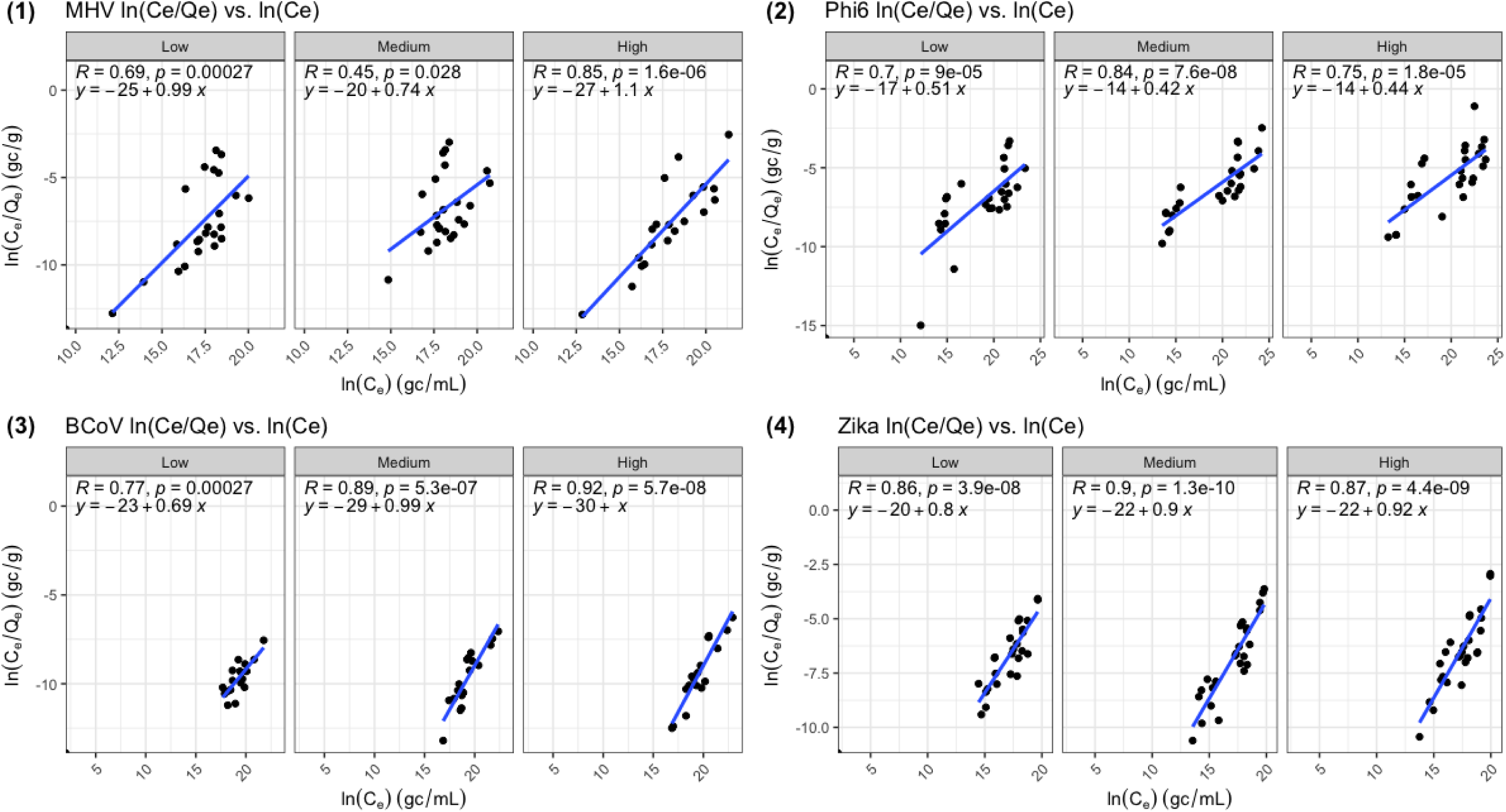
Final Redlich-Peterson isotherm plot for (1) MHV; (2) Φ6; (3) BCoV; (4) ZIKV by TSS. R = r^2^.

**Table 2.** Langmuir, Freundlich, Redlich-Peterson isotherm constants and coefficients by TSS and overall.

|  | **Langmuir** |  |  | **Freundlich** |  |  | **Redlich-Peterson** |  |  |
| --- | --- | --- | --- | --- | --- | --- | --- | --- | --- |
| **System** | $K_L$ (l/g) | $a_L$ (l/mg) | $r^2$ | $K_F$ (l/g) | $n$ | $r^2$ | $K_R$ (l/g) | beta | $r^2$ |
| **BCoV - Low** | $-1.31 \times 10^3$ | $-1.05 \times 10^{-09}$ | 0.14 | $3.58 \times 10^9$ | 2.86 | 0.50 | $2.79 \times 10^{10}$ | 0.65 | 0.73 |
| **BCoV - Medium** | $2.17 \times 10^4$ | $3.48 \times 10^{-09}$ | 0.88 | $3.93 \times 10^{12}$ | -277 | -0.0065 | $2.54 \times 10^{13}$ | 1.0 | 0.87 |
| **BCoV - High** | $1.08 \times 10^4$ | $1.85 \times 10^{-09}$ | 0.72 | $1.07 \times 10^{13}$ | -20.4 | -0.10 | $9.36 \times 10^{-14}$ | 1.0 | 0.91 |
| **MHV - Low** | $1.85 \times 10^2$ | $-8.33 \times 10^{13}$ | -0.30 | $7.20 \times 10^{10}$ | -41.7 | -0.023 | $1.39 \times 10^{-11}$ | 1.0 | 0.69 |
| **MHV - Medium** | 71.4 | $-3.29 \times 10^{-10}$ | 0.022 | $3.58 \times 10^9$ | 6.67 | 0.091 | $2.79 \times 10^{-10}$ | 0.85 | 0.46 |
| **MHV - High** | -76.9 | $-9.23 \times 10^{-10}$ | 0.16 | $5.32 \times 10^{11}$ | -16.1 | -0.096 | $1.88 \times 10^{-12}$ | 1.1 | 0.85 |
| **Φ6 - Low** | $-3.03 \times 10^3$ | $-1.52 \times 10^{-08}$ | 0.55 | $2.42 \times 10^7$ | 2.00 | 0.66 | $4.14 \times 10^{-08}$ | 0.5 | 0.66 |
| **Φ6 - Medium** | 4.76 | $-9.52 \times 10^{-10}$ | -0.43 | $4.42 \times 10^5$ | 1.54 | 0.91 | $2.26 \times 10^{-06}$ | 0.35 | 0.77 |
| **Φ6 - High** | 90.9 | $1.00 \times 10^{-10}$ | 0.077 | $1.20 \times 10^6$ | 1.69 | 0.82 | $8.32 \times 10^{-07}$ | 0.41 | 0.7 |
| **ZIKV - Low** | $1.75 \times 10^3$ | $4.74 \times 10^{-08}$ | 0.67 | $1.78 \times 10^8$ | 3.57 | 0.49 | $5.60 \times 10^{-09}$ | 0.72 | 0.83 |
| **ZIKV - Medium** | $-5.26 \times 10^3$ | $-2.32 \times 10^{-07}$ | 0.84 | $1.31 \times 10^9$ | 5.26 | 0.38 | $7.58 \times 10^{-10}$ | 0.81 | 0.87 |
| **ZIKV - High** | $1.35 \times 10^3$ | $3.65 \times 10^{-08}$ | 0.61 | $4.85 \times 10^8$ | 4.76 | 0.38 | $2.06 \times 10^{-09}$ | 0.79 | 0.83 |
| **Overall** |  |  |  |  |  |  |  |  |  |
| **BCoV** | -4545 | $-1.36 \times 10^{-09}$ | 0.16 | $1.45 \times 10^{12}$ | 22.7 | 0.079 | $6.91 \times 10^{-13}$ | 0.96 | 0.89 |
| **MHV** | 476.2 | $-6.67 \times 10^{-10}$ | -0.01 | $7.20 \times 10^{10}$ | $2.63 \times 10^4$ | 0.000036 | $1.39 \times 10^{-11}$ | 1 | 0.70 |
| **Φ6** | -434.8 | $1.04 \times 10^{-08}$ | -0.14 | $3.27 \times 10^7$ | 1.75 | 0.78 | $3.06 \times 10^{-07}$ | 0.43 | 0.75 |
| **ZIKV** | 2857 | $9.43 \times 10^{-08}$ | 0.71 | $4.85 \times 10^8$ | 4.55 | 0.41 | $2.06 \times 10^{-09}$ | 0.78 | 0.88 |

Figures 3 and 4 show observed time-dependent adsorption as the average log_10_ gene copies per gram of swab material of each target over a 48 hour (2,880 minute) time period and by TSS. Results suggest that MHV, Φ6, BCoV, and ZIKV adsorption behavior is characterized by increased gene copy concentrations on swab material over 12 hours (720 minutes) sampled.

**Fig. 3.**
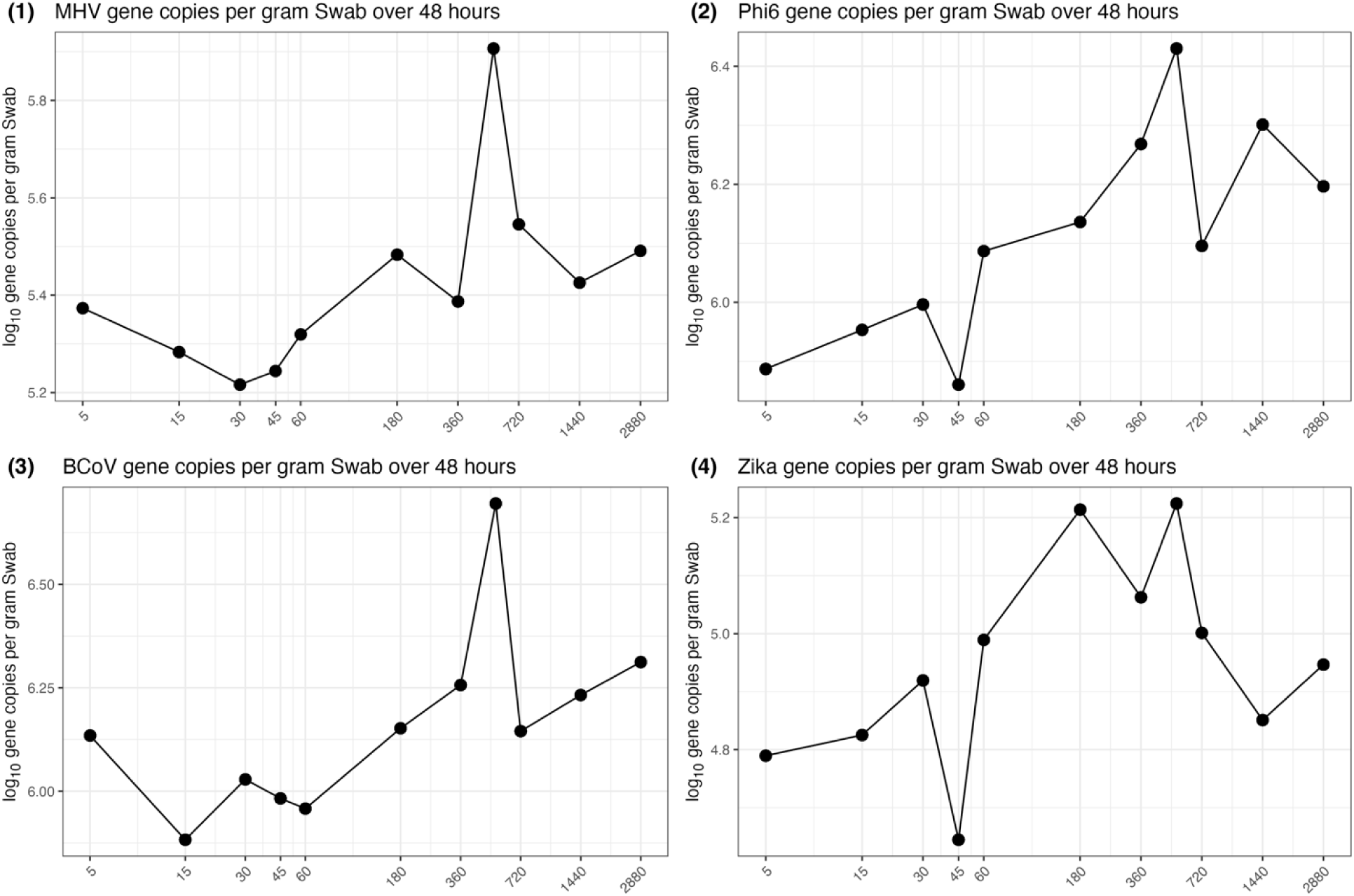
Mean gene copies of (1) MHV; (2) Φ6; (3) BCoV; (4) ZIKV per gram of Moore swab material over 48 hours.

**Fig. 4.**
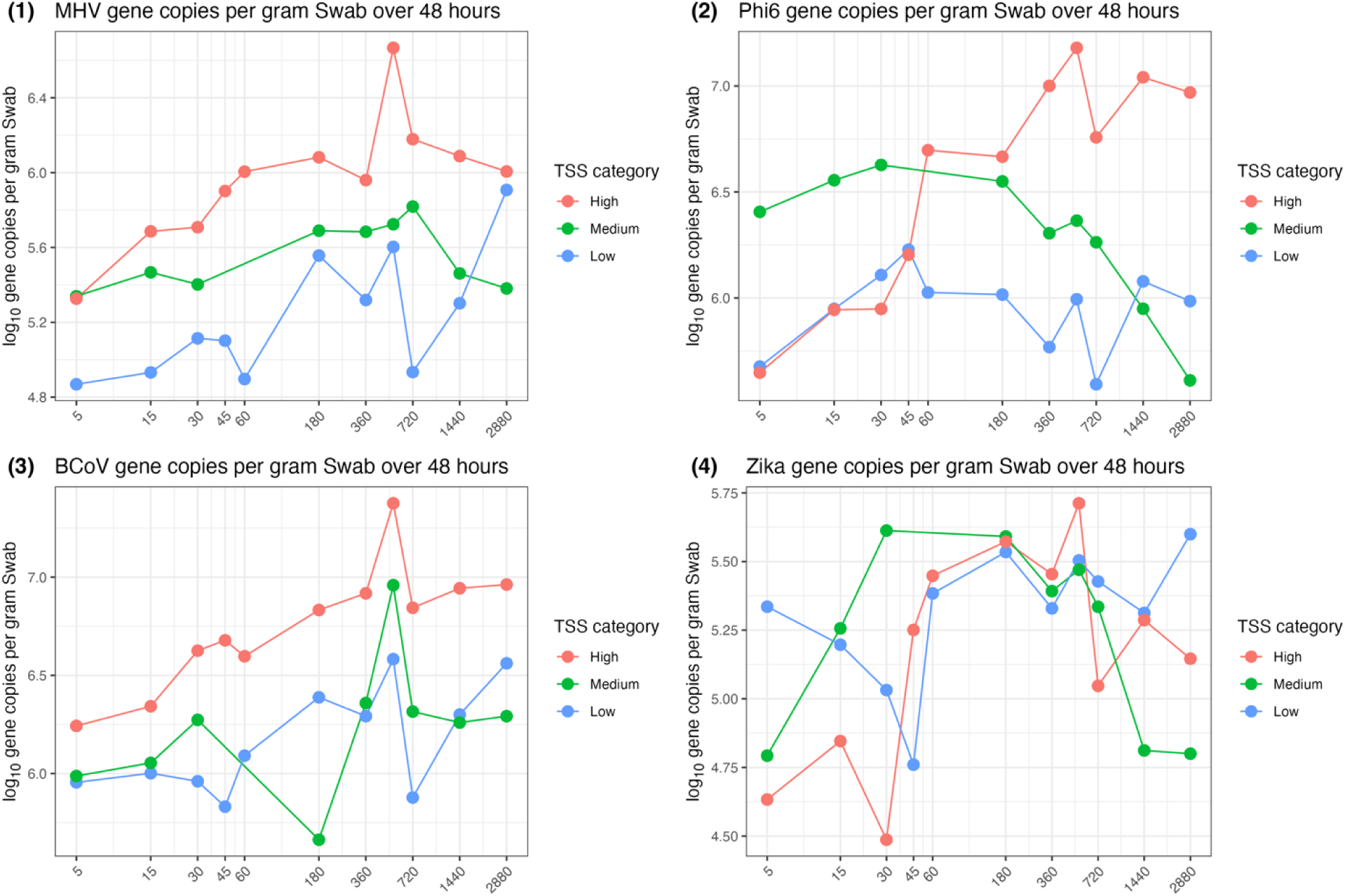
Mean gene copies of (1) MHV; (2) Φ6; (3) BCoV; (4) ZIKV per gram of Moore swab material over 48 hours stratified by TSS categories.

### Desorption

Desorption data (Fig. 5) displays the viral target trend over 48 hours, with the initial concentration plotted as an average of three samples and the final 48^th^ hour concentration performed in duplicate for replicability of initial and final measurements. Although final concentrations were lower than initial concentrations when we placed saturated Moore swabs into initially target-free waters, the rate of desorption was variable, lacking a consistent time-dependent trend. The possibility of a lower viral concentration after 48 hours may also result from degradation of the target.

**Fig. 5.**
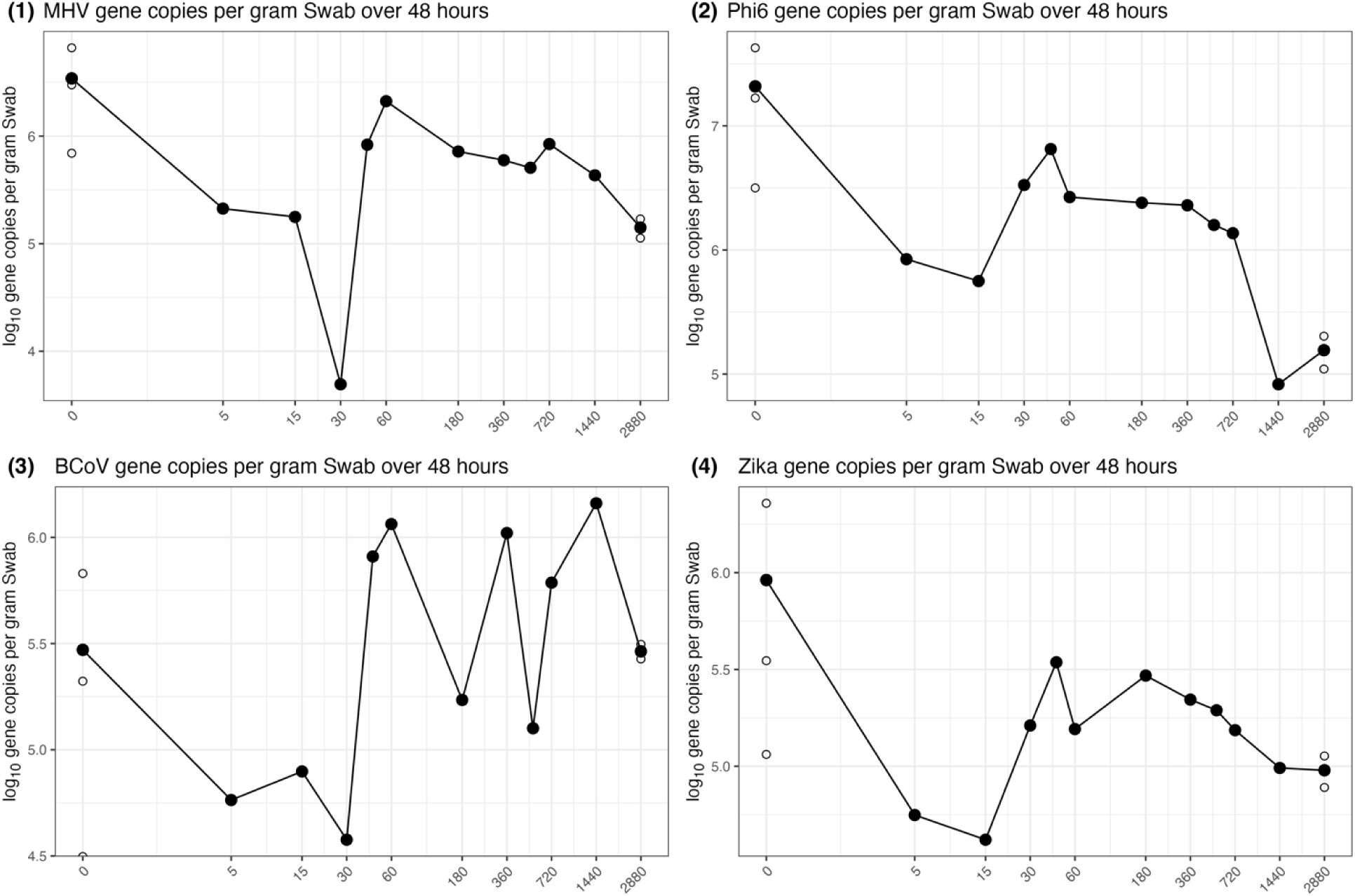
Desorption—Log_10_ gene copies of (1) MHV; (2) Φ6; (3) BCoV; (4) ZIKV per gram of Moore swab over 48 hours.

## Discussion

### Kinetics by target and by TSS

The purpose of the mass loading experiments was to determine the amount of solids and viral targets as gene copies that adsorb to Moore swabs as a function of viral target concentrations in the sampled matrix, which we then translated using isotherm models. The Langmuir models showed poor fit to the data and resulted in a correspondingly low coefficient of determination (R^2^) values for targets, whereas the Freundlich models demonstrated similarly low R^2^ values. However, the Redlich-Peterson model fit performed the best with the highest R^2^ for targets.

### Isotherm relationships

The isotherm relationships can explain how mass loading can be modeled without necessarily testing for concentrations and solids content at every time point. Langmuir isotherms function under specific assumptions, such as adsorption occurs in monolayer in that a single layer of adsorbate molecules occupies the surface. Additional assumptions include that the adsorption surface is homogenous with uniform adsorption sites and that equilibrium adsorption process does not consider kinetic or dynamic processes (Kalam et al., 2021). Redlich-Peterson isotherm models assume a multi-layer adsorption such that multiple layers of adsorbate concentrations can occupy the surface, which can also have multiple adsorption surface sites. The R-P isotherms may also be compatible under a larger adsorbate concentration range, particularly compared to Langmuir isotherm models (Kalam et al., 2021).

Isotherm models describe an empirical relationship whose mechanisms may be highly variable and target– and matrix-dependent. Specific mechanisms may include electrostatic attraction as a function of the viral particle’s isoelectric point (IEP), the pH at which a particle has no net charge, and can play an influential role on viral adsorption to charged surfaces. Viral surfaces exhibit charge differences based on the IEP and the pH of the environment. In a matrix like wastewater, perhaps these mechanisms are negligible, but IEP may still play a role. The IEP for the targets we used in the experiments are as follows: BCoV (4.45-4.65) (Kapil et al., 1999), MHV (6-7.5) (Oleszak et al., 1992), Φ6 (∼6.94) (Moresco et al., 2022), and ZIKV (6.7-7.3) (Poveda-Cuevas et al., 2018). Other mechanisms may include Van der Waals forces, which is the weak intermolecular attraction between all molecules, but neither this nor the IEP can be fully determined or explained through these experiments (Kendall and Roberts, 2015). The isotherms we have derived describe only the empirical loading potential of viruses to surfaces, not the mechanisms by which this is achieved. And, because most viruses are particle-associated in wastewater (Hejkal et al., 1981), sorption behavior of viral RNA is most likely largely determined by the sorption of suspended particles to Moore swab surfaces.

We tested various isotherm fits to determine what model best describes the Moore swab adsorption data and to determine if the underlying, inferred mechanisms could explain adsorption behavior. The R-P model describes complex adsorption, which fits conceptually to the use of Moore swabs in wastewater. Others have considered similar work on electronegative membrane filters and granular activated carbon (GAC) (Hayes et al., 2022a, 2022b), both being best described using the Freundlich and a hybrid Langmuir-Freundlich model, respectively. But no current studies have yet attempted fitting equilibrium and kinetic sorption models to Moore swabs. Our findings suggest that Moore swabs have complex adsorption behaviors, possibly due to the cotton gauze having multiple adsorption sites for targets to attach and detach within a controlled matrix. These results can save time and resources, particularly in knowing that Moore swabs can be used for up to 48 hours as a passive sampler. Additionally, Moore swabs can be readily made in low-resource settings and may prove to be a globally accessible tool. Moore swabs have been used for pathogen detections in other settings such as university surveillance of SARS-CoV-2 in wastewater (Cha et al., 2023; Liu et al., 2022), and gaining larger recognition in passive sampling efforts for wastewater (Bivins et al., 2022a).

### Real world implications

Based on model simulations using the batch equilibrium, if a typical concentration of a target of interest is ∼10^4^ gene copies per mL in the waste stream to be sampled, and, based on our results, a Moore swab made of cotton gauze and ∼5 grams is deployed for 24 hours would have an 85-93% probability of detecting the target, assuming our sample workflow^1^. Given the assay limits of detection, three positive partitions, we suggest using the traditional mass of Moore swabs at ∼ 5g per swab, which generally eluted the highest concentration of targets based on equilibrium experiments. Our experiments demonstrated that if the target concentration of interest is approximately 10^6^ gene copies per mL, then Moore swab deployment for 48 hours can retain a detectable signal. However, we observed the maximum viral target loading at ∼9 hours following deployment, if the viral titer in the water contacting the swab is constant, suggesting that swab deployment during a single day is possible. Conversely, we would expect after 48 hours to lose a positive detection, but based on our scour experiments we were still able to detect all our targets even at 48 hours following removal of the loaded swabs from spiked wastewater. However, we saw a consistent decrease in targets within the first 15 minutes and then a stabilization of concentration between 30-45 minutes (Figure 5). This could indicate our desorption data highlights how well a Moore swab can retain target concentrations after 30 minutes. Other researchers investigating Moore swab deployments recommended a six-hour optimal deployment duration (Cha et al., 2024; Jones et al., 2022), somewhat consistent with the results reported herein.

### Limitations of Moore Swabs

Despite advantages of worldwide low-cost availability, simplicity, and deployability, there are limitations to using Moore swabs. Pre-processing methods for Moore swabs may require more effort than electronegative membranes (Habtewold et al., 2022), but generally yield more grams of solids (i.e., pellet, where targets are enriched by orders of magnitude relative to clarified liquid (Graham et al., 2021; Kim et al., 2022) than a duplicate membrane. Equipment needs for the bead-beating step for electronegative membranes can also be an additional cost burden, especially for those in low-resource settings. Greater recovery of solids may allow for multi-parallel detection and quantification methods that use lower analytical volumes and therefore benefit from relatively high target concentrations, such as those in wastewater (Rao et al., 2024). The explicit combination between target, extraction method and collection device used to optimize pathogen detection still require considerable research. In addition to the fact that we assessed genomic target material and we may not have intact viruses as our endpoint product. There are distinct advantages to using electronegative membranes (Habtewold et al., 2022). Moore swabs have already been criticized for having difficult signal-to-noise ratios (Hayes et al., 2022b) and we can agree that Moore swab experimental data across equilibrium, kinetic, and desorption experiments had unclear trends when samples were averaged together, but when stratified by TSS we did not observe distinct relationships. However, we did not include a clear step to remove inhibitors that may have concentrated through the sample processing methods. Even though we used a sensitive molecular detection platform, there is a possibility of inhibition occurrence. Wastewater is a complex, heterogeneous matrix (Choi et al., 2018) and while we performed all the experiments under controlled conditions, it is possible using different solids concentrations could yield different findings. We consider reported Redlich-Petersen isotherm models to reasonably represent viral loading under the conditions we examined. The high retention of viral targets to Moore swabs over time is an advantage for applications where target presence in the wastewater is variable or fleeting. Further, they could be used for quantitative estimation if concentrations of the target in the wastewater to be sampled is stable, though they should not be considered a substitute for composite sampling or autosampling methods where that is possible. Because of the high degree of uncertainty for any individual quantitative estimate deriving from target recovery via Moore swabs, the most promising application might be in comparing relative concentrations over time in longitudinal sampling of wastewater if flow, wastewater characteristics (e.g., TSS), and time deployed are comparable in repeat sampling events. Deployed in this way, Moore swabs could yield important information about the directionality of target trends over time, which is among the most useful applications of wastewater surveillance (Bivins et al., 2022a; Cha et al., 2024, 2023).

## Conclusions

Viral adsorption on Moore swabs can be characterized under controlled conditions. The Redlich-Peterson isotherm model provides a reasonable fit for Φ6, MHV, BCoV, and ZIKV viral loading on Moore swabs. We find that Moore swab data can be used to generate quantitative estimates, though results should be considered approximate at best and only apply when concentrations of the target in wastewater are stable. We further find that deployment for 9 – 12 hours results in peak loading onto the Moore swab, so single-day sampling events are acceptable if the goal is to maximize opportunity for detection of targets and speed up the time from deployment to results. Our results complement other work characterizing the application of passive sampling methods for wastewater surveillance as a scalable method particularly suited to underserved settings where autosamplers may not be available.

## CRediT authorshop contribution statement

Gouthami Rao: Writing – original draft, Visualization, Methodology, Formal analysis, Data curation. Timothy Purvis: Investigation, Methodology, Writing – review & editing. Gyuhyon Cha: Validation, Writing – review & editing. Jack Dalton: Methodology, Data curation, Writing – review & editing. Michael Fisher: Methodology, Writing – review & editing. Katherine Graham: Validation, Writing – review & editing. Konstantinos Konstantinidis: Project administration, Writing – review & editing. Yarrow Linden: Methodology, Writing – review & editing. Aaron Bivins: Validation, Writing – review & editing. Margo A. Brinton: Resources, Writing – review & editing. Christine Stauber: Methodology, Funding acquisition, Writing – review & editing. Joe Brown: Writing – review & editing, Methodology, Conceptualization, Supervision, Funding acquisition.

## Acknowledgements

This study was funded by the National Science Foundation (NSF Award #2027758). The funders had no role in study design, data collection and analysis, decision to publish, or preparation of the manuscript. Further we would like to acknowledge members of the Stauber, Fisher, and Brinton labs for the technical culturing efforts.

## Footnotes

1 The probability of detecting a target was calculated by fitting a probit model to the observed equilibrium data and simulated detection probabilities (n=1000) at lower concentrations (0.1, 0.5, 1 gene copies/mL).

